# MicroBayesAge: A Maximum Likelihood Approach to Predict Epigenetic Age Using Microarray Data

**DOI:** 10.1101/2024.10.04.616756

**Authors:** Nicole Nolan, Megan Mitchell, Lajoyce Mboning, Louis-S. Bouchard, Matteo Pellegrini

## Abstract

Certain epigenetic modifications, such as the methylation of CpG sites, can serve as biomarkers for chronological age. Previously, we introduced our BayesAge frameworks for accurate age prediction through the use of locally weighted scatterplot smoothing (LOWESS) to capture the non-linear relationship between methylation or gene expression and age, and Maximum Likelihood Estimation (MLE) for bulk bisulfite and RNA sequencing data. Here we now introduce MicroBayesAge, a framework that enhances prediction accuracy by subdividing input data into age-specific cohorts and employing a new two-stage process for training and testing. Age prediction for younger patients was significantly improved. MicroBayesAge also exhibited minimal bias in its age predictions. Additionally, we explored the performance of our model for sex-specific age prediction which revealed slight improvements in accuracy for male patients, while no changes were observed for female patients. MicroBayesAge provides more accurate age predictions by accounting for variations in epigenetic markers of aging among different subgroups, which have been overlooked by commonly used models.

## 1 INTRODUCTION

While human genetic sequences remain broadly unchanged over time, the epigenome is highly dynamic and subject to modifications that can be influenced by environmental factors, lifestyle, and aging. Epigenetic modifications to DNA occur over the course of a lifetime, and certain epigenetic changes can be used as a means of determining a person’s age [1]. These modifications include histone modifications, non-coding RNA interactions, and, most prominently, DNA methylation [2]. In particular, CpG site methylation, in which a methyl group is added to the C5 of the cytosine in a CpG dinucleotide, can serve as a robust epigenetic indicator for aging [3]. This process affects gene expression without altering the underlying genetic code and can be influenced by both intrinsic and extrinsic factors. CpG methylation patterns have been widely studied and linked to various biological processes, including cellular differentiation, senescence, and disease progression [4]. As a result, DNA methylation at specific CpG sites has become a reliable biomarker for estimating biological age, a concept that has implications for aging research, forensic science, and personalized medicine.

Different methods of predicting chronological age have been developed, commonly referred to as “epigenetic clocks.” [5–8]. Several of these epigenetic clocks involve DNA methylation analysis using various types of regression algorithms, such as elastic net regression or penalized regression. The development of these clocks has provided valuable tools for aging research; however, the accuracy and applicability of the predictions can vary depending on the population, the tissue type analyzed, and the specific algorithm used. Thus, there is ongoing research to refine these models to improve their precision, reduce biases, and increase their generalizability across different cohorts.

Very few methods developed accounted for the non-linearity trends between methylation and age [8,9]. Previous research has shown that, for many CpG sites, methylation changes occur more rapidly at younger ages and more slowly at older ages, suggesting a relationship that is more logarithmic than linear [10]. Recognizing the limitations of existing linear and penalized regression models, we recently developed our own age prediction algorithm, BayesAge 1.0 [11], which predicts age from CpG site methylation data using a combination of locally weighted scatterplot smoothing (LOWESS) regression and Maximum Likelihood Estimation (MLE). BayesAge 1.0 used LOWESS smoothing to account for this nonlinear relationship, resulting in less biased age predictions when compared with common epigenetic clocks which assume a linear relationship. For BayesAge 2.0, we implemented a similar framework as it’s predecessor to predict transcriptomic age based on a gene expression profiles [12]. However, while BayesAge 1.0 and 2.0 was developed to predict the epigenetic and transcriptomic age from bulk bisulfite and RNA sequencing datasets, they are not applicable to predicting biological age from microarray technology such as the Illumina 450K BeadChip platform [13].

Herein, we introduce MicroBayesAge, a bayesian method for epigenetic age prediction utilizing microarray data. This method leverages the most highly correlated CpG sites, identified via Spearman rank correlation, for age prediction. During the training phase, LOWESS smoothing captures the non-linear relationship between methylation and age for each selected CpG site, and variance per CpG site is calculated to create reference matrices for predefined ages. In the prediction phase, the mean and variance are used to fit a normal distribution to each top CpG site. The sum of these top correlated CpG sites is used to generate a maximum likelihood age distribution, providing an accurate estimate of biological age. MicroBayesAge offers a streamlined and precise method for epigenetic age prediction to enhance our understanding of the aging process. This innovative approach improves upon previous models by offering robustness to missing data and increased interpretability. Furthermore, we implement a two-stage Bayesian framework that predicts age for young patients extremely accurately, within around 2.5 years of the chronological age prediction. Finally, we explore the prediction accuracy of our models when the training dataset is divided by sex and find that the prediction improvements are minimal if at all present.

## 2 MATERIALS AND METHODS

### 2.1 Data Sourcing and Analysis

This version of the BayesAge framework is designed to predict the age of a person based on CpG site methylation data collected from their DNA. To develop and test MicroBayesAge, we utilized a dataset downloaded from the NCBI Gene Expression Omnibus [14]. This dataset comprises publicly available methylome data derived from human blood samples gathered in 19 separate epigenetic studies [15–33]. The data were obtained using the Illumina HumanMethylation450 BeadChip for genome-wide CpG site methylation analysis of DNA samples.

In total, the dataset contains methylomes from over 11,000 patients, each with methylation data for over 200,000 CpG sites, along with their chronological age and sex. These patients span a wide range of ages, from newborns to centenarians (see Fig. 1), providing a broad sample base for training and validation of the algorithm.

**FIGURE 1.**
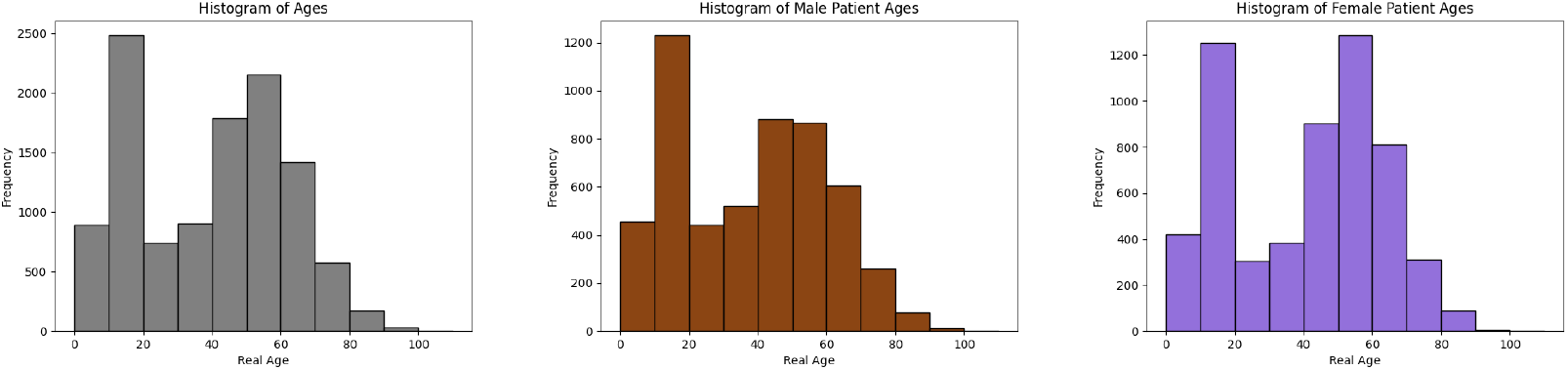
Distribution of real ages for all patients, shown in gray, and for male and female patients in sex-specific datasets, shown in brown and purple respectively.

A Spearman’s rank correlation coefficient between patient age and methylation was calculated for each of the CpG sites. Spearman correlation was used because it is robust for nonlinear trends, such as the logarithmic relationship between methylation and age. We removed from consideration all CpG sites whose Spearman rank correlation had an absolute value of less than 0.60 to focus on the methylation data most relevant for age prediction. This process narrowed the number of CpG sites under consideration to approximately 1,500.

To evaluate the MicroBayesAge framework, we employed ten-fold cross-validation. The dataset was split into two groups of samples: testing groups and a training groups. The testing groups were created by subdividing the full dataset into ten mutually exclusive subsets of randomly selected samples, each comprising 10% of the total number of samples. This generated random groups of approximately 1,000 test samples from the overall dataset of over 11,000 samples. For each testing group, the remaining 90% of the samples from the full dataset were used as the corresponding training group.

### 2.2 MLE Algorithm

The MLE algorithm in MicroBayesAge functions similarly to that of BayesAge 1.0, but it has been updated to handle microarray data. BayesAge 1.0 utilized targeted bisulfite sequencing (TBS-seq) to obtain direct counts of methylated and unmethylated cytosines at each CpG site. It was proposed that the probability of measuring the observed counts, given the expected methylation level for a particular age, followed a binomial distribution. The probability for a specific CpG site state could then be computed as a function of the number of reads of cytosines and the number of total reads.

MicroBayesAge, on the other hand, is trained on microarray data from the Illumina HumanMethylation450 Bead-Chip. In place of counts, signal intensities are converted into a single beta value that represents the methylation level for a particular CpG site [34]. Without count data, we cannot model the probability as a binomial distribution. Instead, we use the expected methylation levels from the trained model along with variance to model the probability distribution for a distinct age.

Considering that variance may change at differential levels of methylation [35], we chose to estimate variance by treating samples of the same age as technical replicates. We then plotted the mean methylation level within each age bin against its variance and used LOWESS smoothing to produce a unique curve of methylation vs. variance for each CpG site. We tested *τ* values from 0.1 to 1 in intervals of 0.1 and determined that a value of 0.7 resulted in predictions with the lowest MAEs. For each expected methylation level from the reference matrix, we sampled the predicted variance from this curve and calculated its square root to estimate the standard deviation for a specific age and methylation level.

We then tested both a beta distribution and normal distribution to compute the posterior probability. To fit a beta distribution, we used the mean methylation level from the reference matrix and the variance of a given CpG site to calculate the shape parameters for a distinct age as follows:

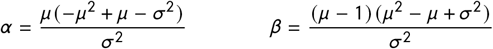

where:

*α, β* : Shape parameters.

*µ*: Mean methylation level for a given age.

*σ*^2^: Variance for given age and methylation level.

The posterior probability of a given measurement for a particular age was then computed using the `beta.pdf` function of the `scipy.stats` package. The associated probability for a unique CpG site state is described using the formula below, where 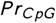 represents the probability for a given methylation level at a particular age:

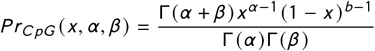

where:

Γ is the gamma function. (scipy.special.gamma)

*x* : Measured methylation level for this subject.

*α, β* : Shape parameters calculated above.

However, we found that our model performed no differently when calculating the posterior probability using a normal distribution. For simplicity, the posterior probability was modeled using the `norm.pdf` function of the `scipy.stats`, as described below [36].

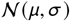

where:

*µ*: Expected methylation level at the given age.

*σ*: Standard deviation for this CpG site at the given age.

To calculate the probability of observing the methylation level measured across all CpG sites for a particular age, we take the logarithmic sum of the probability for each of the top 16 sites. In this manner the algorithm calculates the likelihood of observing each age from 0 to 100 in a single subject, producing an age likelihood distribution. The maximum likelihood age is predicted as the epigenetic age of the subject.

### 2.3 Two-Stage MicroBayesAge Framework

The original BayesAge 1.0 framework consisted of two phases: a training phase and an age prediction phase. In the training phase, a small subset of CpG sites exhibiting the highest absolute value Spearman rank correlation with age was selected for analysis. For MicroBayesAge, extensive optimization testing was performed to determine the optimal number of CpG sites to be used in the training phase to achieve the highest prediction accuracy. After the selection of CpG sites was made, LOWESS regression was applied to model the relationship between age and the methylation levels of the selected CpG sites. We used the `lowess` function from version 0.14.1 of the `statsmodels.api` Python module [37]. Following further optimization testing, we determined the optimal tau parameter for the smoothness of the LOWESS fit.

In developing MicroBayesAge, we theorized that the relationship between CpG site methylation and age might differ across different age groups and that more accurate age predictions could be achieved if these differing relationships were accounted for. To test this hypothesis, MicroBayesAge utilizes a two-stage training phase and a two-stage prediction phase to improve accuracy (see Fig. 2).

**FIGURE 2.**
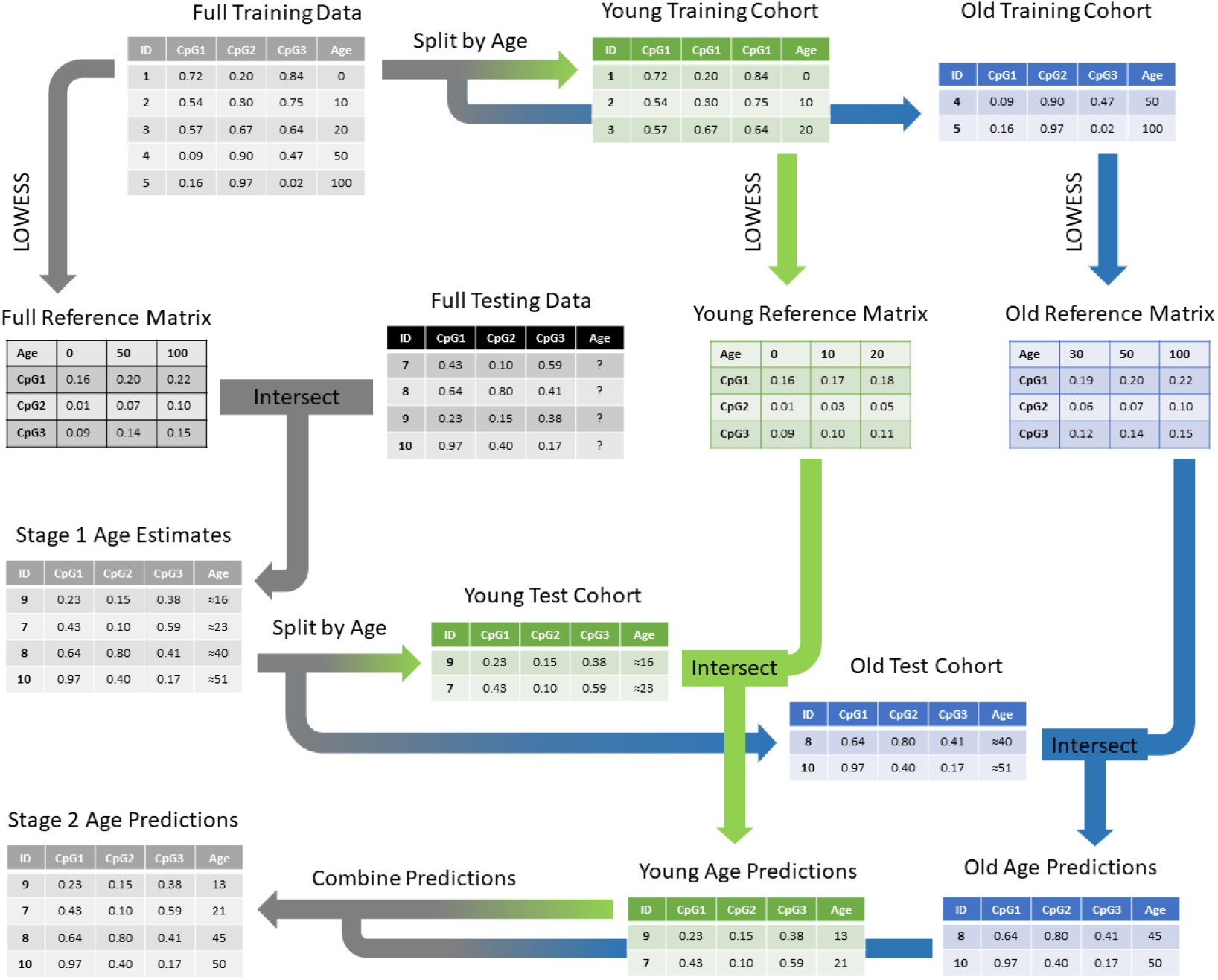
Schematic of the BayesAge version 3.0 framework.

In the first stage of the training phase, MicroBayesAge develops its age prediction algorithm using an initial training dataset consisting of CpG site methylation data from individuals with known ages. The framework calculates the Spearman rank correlation coefficient between real age and methylation for each CpG site, using only the available training dataset. The top 16 CpG sites most strongly correlated with age are then selected based on these Spearman ranks and the `lowess` function is used to compute the fit for each of the 16 sites.

In the second stage of the training phase, the training dataset is split into two groups: samples with real ages of 25 or younger, and samples with real ages of 26 or older. For each of the age cohorts, the Spearman rank correlation is recalculated for all CpG sites using only the data from that age cohort. Based on the newly calculated Spearman ranks, a new set of the top 16 most correlated CpG sites is identified for each age cohort and new LOWESS fits are then calculated for the top 16 CpG sites in each cohort.

Once the training phase is complete, the trained model consists of three reference matrices, each composed of 16 CpG sites and their corresponding methylation levels across possible ages, ranging from 0 to 100 in 1-year increments. With this trained model, MicroBayesAge can then be used to predict ages for samples with known CpG site methylation levels and unknown real ages.

When given a testing dataset with unknown real ages to analyze, a two-stage prediction process is employed to predict the age of each sample. For the first stage of the prediction process, the reference matrix developed in the first stage of the training process is intersected with the corresponding CpG sites in the testing dataset. MicroBayesAge then employs Maximum Likelihood Estimation (MLE) to generate initial age estimates based on the intersection of the reference matrix with the CpG site methylation data from the testing dataset.

In the second stage of prediction, the algorithm subdivides the input datasets into two different age cohorts based on the initial age estimations. Specifically, a young cohort is created, comprising samples with predicted age of 25 or younger, and an old cohort is created, comprising samples with predicted ages of 26 or older. For each of these age cohorts, the corresponding reference matrices generated in the second stage of the training phase are intersected with the CpG site methylation data. New age predictions are then separately calculated for each cohort using MLE.

This two-stage training and prediction process is designed to determine if MicroBayesAge can provide more accurate age predictions by accounting for potential differences in the relationship between CpG site methylation and age for different age cohorts. To determine the relative accuracy of the first-stage predictions, which do not attempt to account for any potential differences between age cohorts, compared to the second-stage predictions, we calculated the mean absolute error (MAE) of our age predictions. For both the first-stage and second-stage age predictions, the MAEs for the age predictions of the entire patient population were calculated, as well as the MAEs for the young age cohort and the old age cohort specifically.

### 2.4 Sex-Specific BayesAge Framework

We further theorized that sex-based differences in the relationship between CpG site methylation and age could be accounted for to increase the accuracy of our age predictions [38]. To test this, we split the entire training dataset into two groups by sex and similarly split the entire testing dataset into two groups by sex. This resulted in male and female training datasets as well as male and female testing datasets. We then trained and tested the MicroBayesAge framework on the male and female training and testing datasets separately. Comparing the MAEs calculated for the sex-specific predictions against the MAEs calculated for the full mixed-sex testing dataset allowed us to determine the impact of sex-specific modeling on the accuracy of age prediction.

### 2.5 LASSO Regression Comparison

To provide a point of comparison against MicroBayesAge, we utilized least absolute shrinkage and selection operator (LASSO) regression, which is commonly employed alongside ridge regression for age prediction in a regression method known as elastic net [39]. For this we used the `Lasso` function of the `scikit-learn` python package. We split our overall dataset into training and testing data for the LASSO regression using the same random selection method employed to train and test BayesAge After extensive optimization testing of parameters ranging from 0.01 to 1.00 in increments of 0.01, a lambda parameter value of 0.01 was found to provide the most accurate age predictions.

## 3 RESULTS

### 3.1 Prediction by Age Cohort

Our objective was to determine if the BayesAge framework could generate more accurate age predictions by examining specific age cohorts of patients individually. For the age-specific training and testing of MicroBayesAge, we developed a two-stage training and prediction process. After optimization testing, we found that utilizing the top 16 most highly correlated CpG sites for age prediction gave optimal results.

In the first stage of training, the top 16 CpG sites showing methylation levels with the highest Spearman correlation with age were selected based on the entire patient dataset (see Fig. 3). A LOWESS regression fit of the relationship between methylation and age for these 16 CpG sites was then calculated. These relationships were subsequently used to formulate the first stage of the age predictions for all samples within the validation dataset.

**FIGURE 3.**
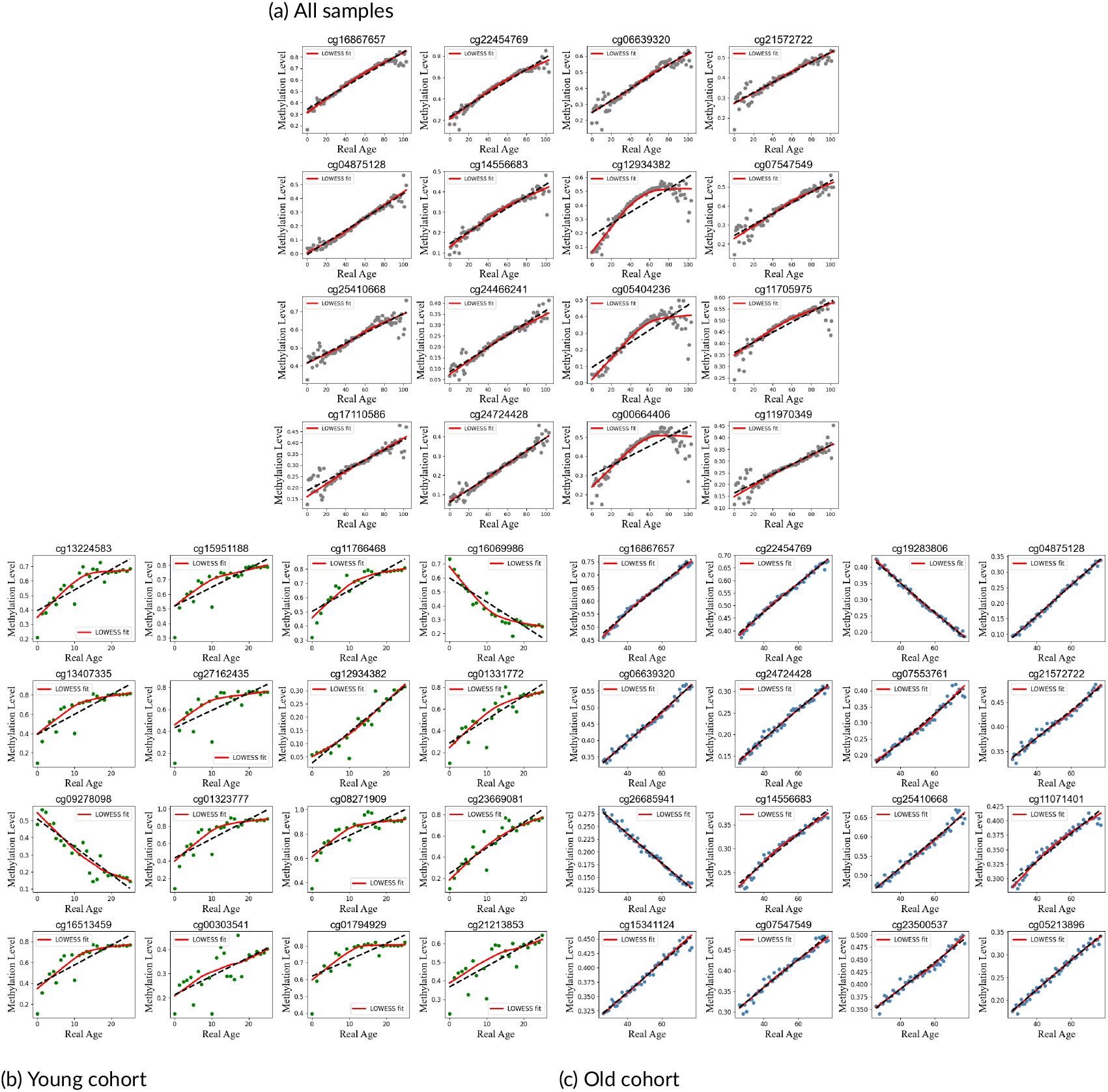
Comparison of LOWESS regression fits with tau of 0.7 of the relationship between methylation and age for the top 16 most correlated CpG sites for all samples, the young cohort only, and the old cohort only.

The top 16 most correlated CpG sites for each of the two age cohorts were then selected, and similar LOWESS regression fits were calculated for both age cohorts (see Fig. 3). Because separate subsets of patients were considered, the 16 CpG sites selected varied for each of the two age cohorts, and new regression fits were calculated. These newly calculated relationships were then used to generate the second stage of age predictions for the young and old samples in the validation dataset separately (see Fig. 4).

**FIGURE 4.**
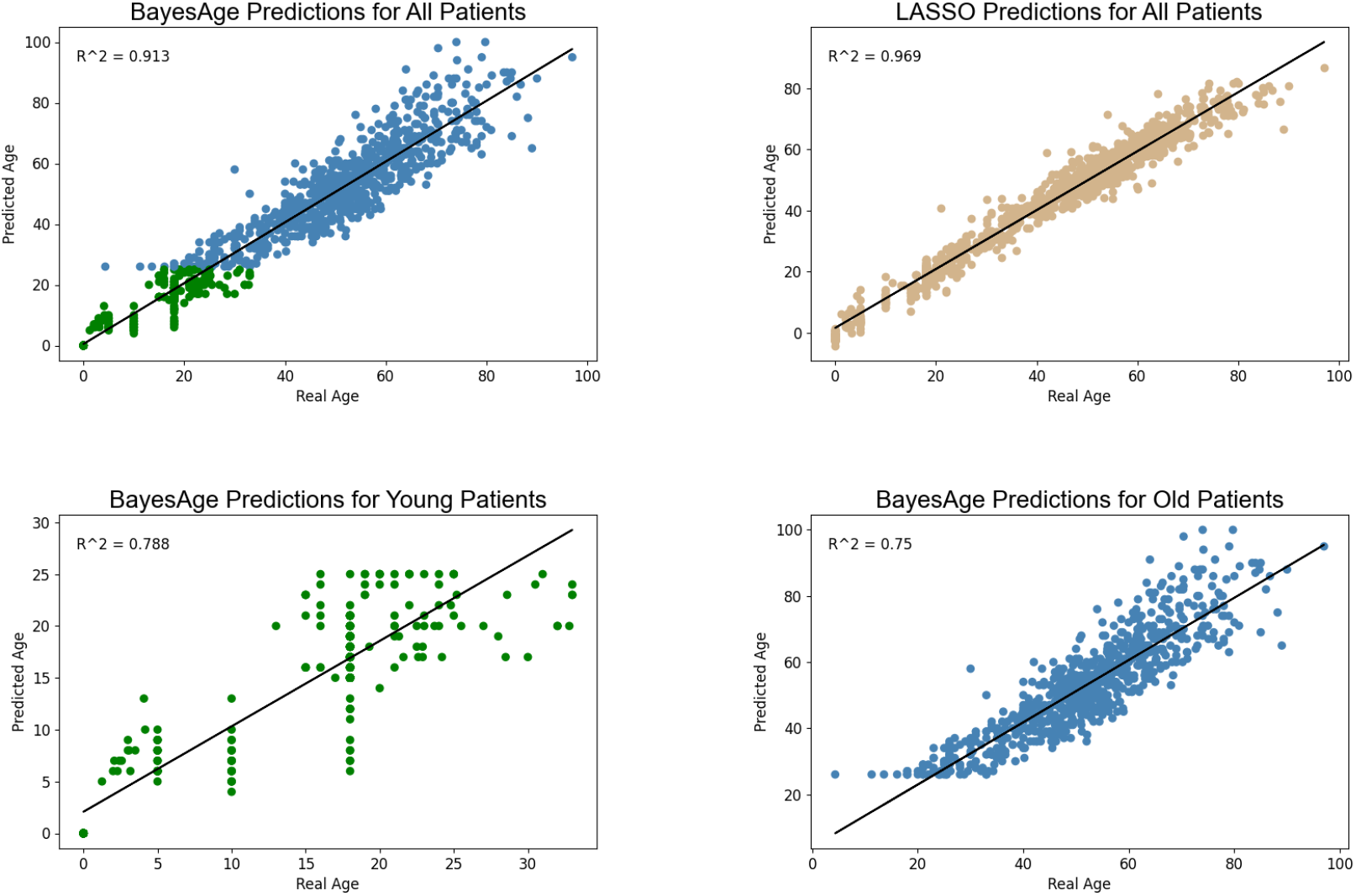
Comparison of second stage age predictions plotted against real age for each age group of patients. Patients with BayesAge predictions older than 25 are shown in blue while patients with predictions of 25 or younger are shown in green. LASSO age predictions are shown in tan for comparison.

In order to determine the bias of our predictions we calculated the residuals, that is the difference between our predicted ages and the real ages, for each of the testing samples. One of the advantages that other versions of BayesAge exhibited over alternate methods of age prediction, such as scAge, were comparatively unbiased age predictions, meaning there was minimal correlation between the residuals and the real age of the samples. Both the first-stage and second-stage predictions of MicroBayesAge similarly exhibit relatively unbiased age predictions when compared to our LASSO benchmark age predictions, although we found no additional reduction in bias for the second-stage predictions relative to the first-stage predictions. (see Fig. 5).

**FIGURE 5.**
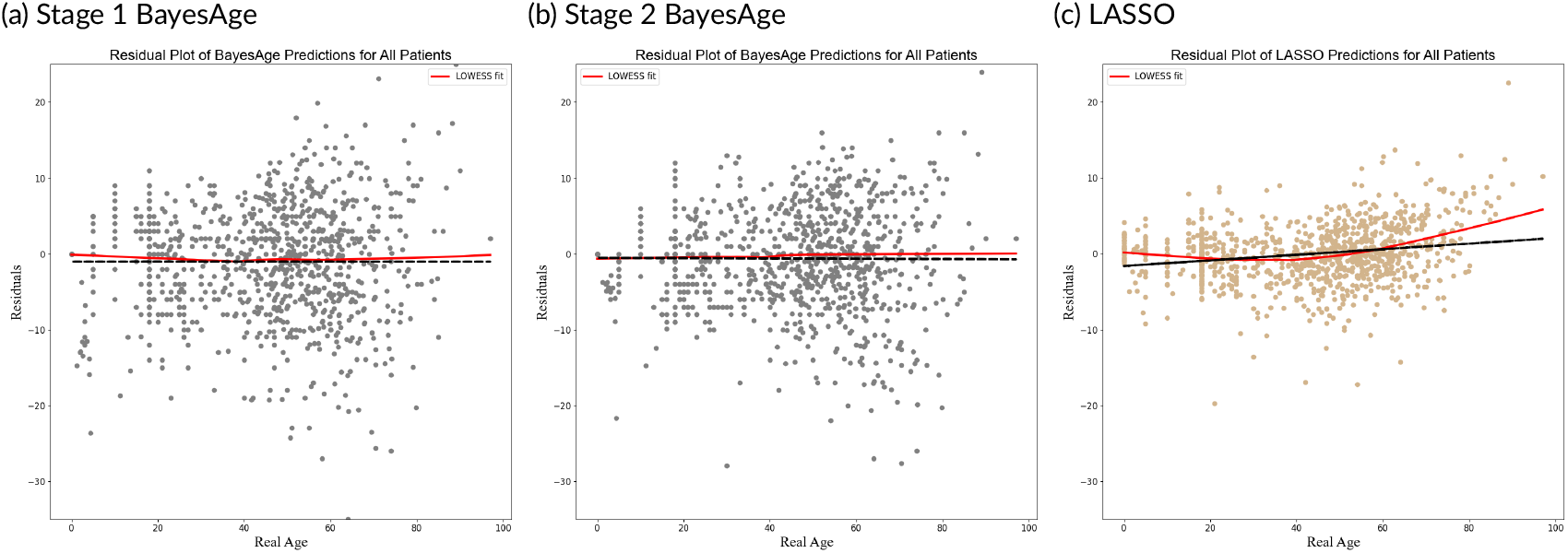
Residual plots of age predictions. First stage age predictions for all patients are shown in gray on the left. Second stage age predictions are shown in gray in the center. LASSO age predictions are shown in tan on the right.

After testing, we found that the addition of the second stage of our age prediction algorithm significantly decreased the Mean Absolute Error (MAE) of the age predictions. This improvement in accuracy was primarily driven by a large decrease in the MAE of age predictions for young patients, defined as those who were 25 years old or younger. There was comparatively less change in the MAE of age predictions for patients older than 25 years, with only a small increase in accuracy observed (see Fig. 6).

**FIGURE 6.**
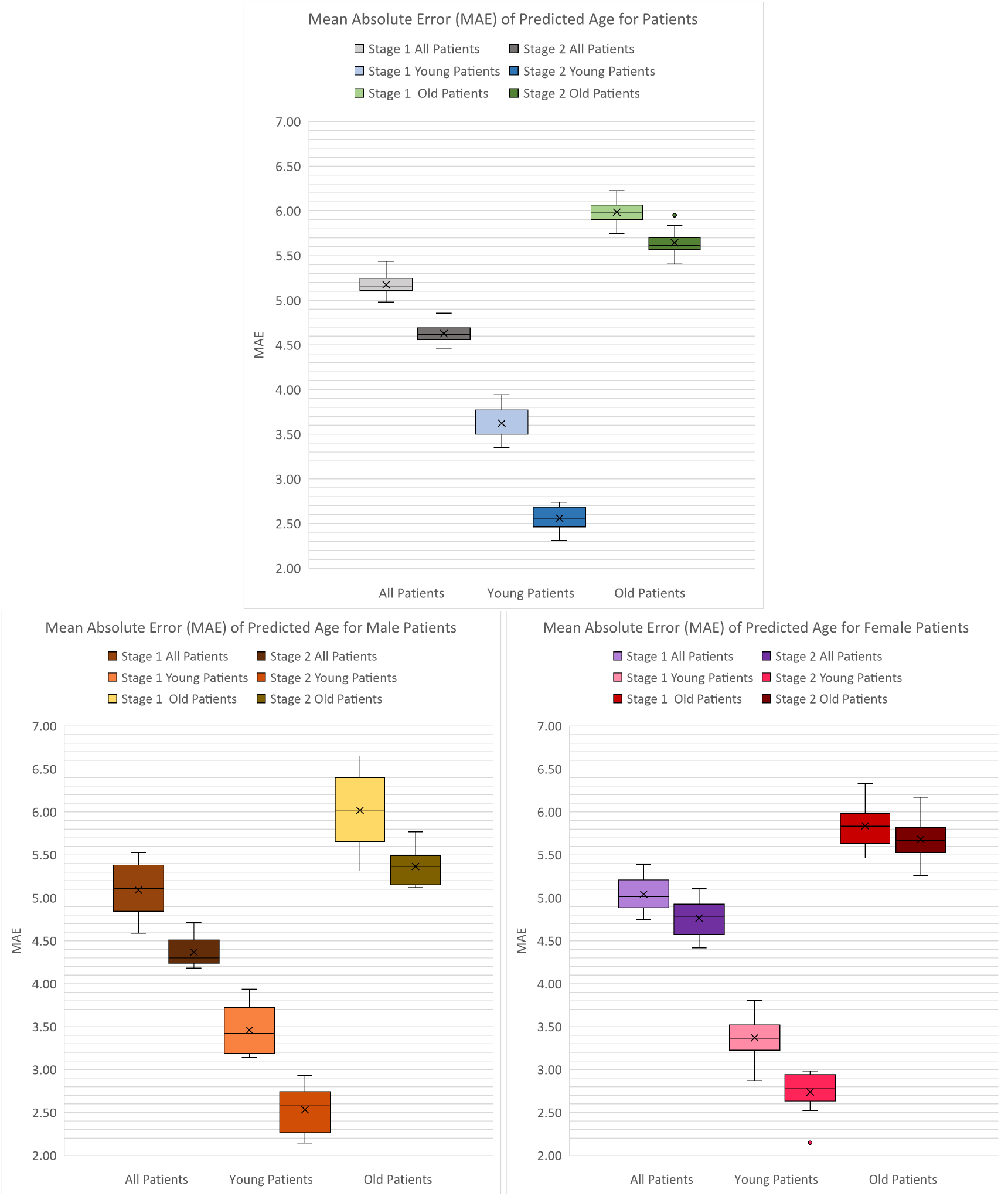
Comparison of MAE for first and second stage age predictions for each age cohort of patients across all patients and across sex-specific datasets.

The overall MAE for patients across all ages improved significantly, decreasing from 5.17±0.04 years for the first-stage predictions to 4.63±0.03 years for the second-stage predictions. This increase in accuracy of the age predictions was primarily due to the improvement in accuracy for patients 25 years old or younger. For this group of younger patients, the MAE decreased from 3.62±0.05 years for the first-stage predictions to 2.56±0.04 years for the second-stage predictions, a substantial decrease. There was comparatively little improvement in the accuracy of age predictions for patients older than 25 years. The MAE for older patients decreased from 5.99±0.05 years for the first-stage predictions to 5.64±0.05 years for the second-stage predictions. For comparison, LASSO regression showed an overall MAE of 2.77±0.03 years for all patients.

### 3.2 Prediction by Sex

The sex-specific datasets, created by splitting the training and testing datasets into male and female groups, each contained a broad range of ages distributed similarly to those of the original full dataset (see Fig. 1).

With the sex-specific datasets, slight increases in the overall accuracy of age predictions were found for male patients; however, no significant change was observed in the accuracy of age predictions for female patients compared to the predictions based on the full dataset of all patients (see Fig 6).

For male patients, the overall MAE for patients across all ages improved slightly, decreasing from 4.63±0.03 years for the mixed-sex second-stage predictions to 4.37±0.06 years for the sex-specific second-stage predictions, an improvement primarily due to an increase in age prediction accuracy for older patients. For patients 25 years old or younger, the sex-specific MAE of 2.53±0.09 years was comparable to the mixed-sex MAE of 2.56±0.04 years. For patients older than 25, the sex-specific MAE of 5.36±0.07 years was slightly lower than the mixed-sex MAE of 5.64±0.05 years.

For female patients, the overall MAE for patients across all ages was 4.76±0.07 years, showing no significant change compared to the mixed-sex MAE of 4.63±0.03 years. The MAEs for the sex-specific young and old patient cohorts were similarly comparable for female patients to those of the mixed-sex age predictions.

## 4 DISCUSSION

We developed a new version of our BayesAge framework, which generates more accurate age predictions from the analysis of DNA methylation data than previous iterations. To achieve this, MicroBayesAge leverages the differences in the relationship between methylation and age for younger versus older individuals. Using a two-stage process for both training and prediction, MicroBayesAge is able to account for these variations between age cohorts when constructing prediction algorithms and categorizes data with unknown ages via initial rough estimates before generating more precise predictions. This subdivision of input data into cohorts based on estimated age allows MicroBayesAge to apply specifically tailored analyses to each age cohort individually.

The second stage of MicroBayesAge shows significantly increased overall age prediction accuracy when compared to the first stage. Notably, it achieves much greater accuracy in predicting ages for the younger cohorts, but shows comparatively less dramatic improvements in the accuracy of the age predictions for the older cohorts. The relationship between methylation and real age for the most correlated CpG sites appears to be nearly linear for older patients while younger patients exhibit more nonlinear relationships. This may explain why our BayesAge algorithm, which is designed to better account for nonlinear relationships, produces more accurate predictions for the younger cohort than for the older cohort. In addition, the residuals of the age predictions indicate minimal bias for both the first and the second stage age predictions generated by MicroBayesAge when compared to the predictions produced by the LASSO benchmark.

We also considered the possibility that male and female patients might, similar to the young and old patient cohorts, exhibit differing relationships between age and CpG site methylation. To test this, we split the patients in the training and testing datasets into two groups based on sex. The BayesAge framework was then trained and tested on male and female patients separately to determine if accuracy would improve with sex-specific training and age prediction. We found slight improvements in age prediction accuracy for male patients when compared to the mixed-sex predictions, but no change in accuracy for female patients.

In conclusion, MicroBayesAge uses the partitioning of training datasets and input data to achieve greater accuracy. By separating broad datasets into more specific groups, it can better account for differences in epigenetic aging between different classes of patients. This strategy has great potential for further expansion. Age-based partitioning could be further optimized to generate more refined predictions, and further means of subdividing data based on other criteria could potentially provide further improvements in accuracy. For instance, it could improve accuracy further if patients were subdivided into a greater number of smaller, more focused age cohorts. Additional data points, such as information regarding chronic medical conditions, might also be used to subdivide datasets and potentially produce more accurate predictions.

## Supporting information

MicroBayesAge Supplemental

## 5 CODE AND DATA AVAILABILITY

The code and data used for this study are available at: https://github.com/nicolecnolan/MicroBayesAge

## 6 AUTHOR CONTRIBUTIONS

**NN:** Data curation, Methodology, Software, Validation, Visualization, Writing–original draft, Writing–review and editing. **MM:** Data curation, Methodology, Software, Validation, Visualization, Writing–original draft, Writing–review and editing. **LM:** Conceptualization, Formal Analysis, Investigation, Methodology, Software, Writing–review and editing. **LSB:** Supervision, Writing-review, and editing. **MP:** Conceptualization, Methodology, Supervision, Writing-review, and editing.

## 7 ACKNOWLEDGEMENTS

Nicole Nolan is a NSF UCLA fellow. This work was supported by the NSF-UCLA Quantum Science and Engineering PhD Fellowship. Lajoyce Mboning is a Eugene V. Cota-Robles, Warren Alpert Computational Biology and AI Network and a NSF NRT fellow. This work was supported by the NIH Training Grant in Genomic Analysis and Interpretation T32HG002536. We thank Martin Zhang, Michael Thompson, Mathew Soldano and Stella Gray for the discussions.

## 8 CONFLICT OF INTEREST STATEMENT

The authors declare no competing interests.

## Notes

https://github.com/nicolecnolan/MicroBayesAge

### Competing Interest Statement

The authors have declared no competing interest.

## References

[1] Duan R, Fu Q, Sun Y, Li Q. Epigenetic clock: A promising biomarker and practical tool in aging. Ageing Res Rev. 2022 Nov;81:101743. doi: 10.1016/j.arr.2022.101743. Epub 2022 Oct 4. PMID: 36206857.

[2] Bure IV, Nemtsova MV, Kuznetsova EB. Histone Modifications and Non-Coding RNAs: Mutual Epigenetic Regulation and Role in Pathogenesis. International Journal of Molecular Sciences. 2022 May 22;23(10):5801. doi: 10.3390/ijms23105801. PMID: 35628612; PMCID: PMC9146199.

[3] Moore L, Le T, Fan G. DNA Methylation and Its Basic Function. Neuropsychopharmacology. 2013;38(1):23–38. doi: 10.1038/npp.2012.112.

[4] Zhu X, Chen Z, Shen W, et al. Inflammation, epigenetics, and metabolism converge to cell senescence and ageing: the regulation and intervention. Signal Transduction and Targeted Therapy. 2021;6:245. doi: 10.1038/s41392-021-00646-9.

[5] Horvath S. DNA methylation age of human tissues and cell types. Genome Biol. 2013;14(10):R115. doi: 10.1186/gb-2013-14-10-r115. Erratum in: Genome Biol. 2015 May 13;16:96. doi: 10.1186/s13059-015-0649-6. PMID: 24138928; PMCID: PMC4015143.

[6] Lu AT, Quach A, Wilson JG, Reiner AP, Aviv A, Raj K, Hou L, Baccarelli AA, Li Y, Stewart JD, Whitsel EA, Assimes TL, Ferrucci L, Horvath S. DNA methylation GrimAge strongly predicts lifespan and healthspan. Aging (Albany NY). 2019 Jan 21;11(2):303–327. doi: 10.18632/aging.101684. PMID: 30669119; PMCID: PMC6366976.

[7] Farrell C, Snir S, Pellegrini M. The Epigenetic Pacemaker: modeling epigenetic states under an evolutionary framework. Bioinformatics. 2020 Nov 1;36(17):4662–4663. doi: 10.1093/bioinformatics/btaa585. PMID: 32573701; PMCID: PMC7750963.

[8] Galkin F, Mamoshina P, Kochetov K, Sidorenko D, Zhavoronkov A. DeepMAge: A Methylation Aging Clock Developed with Deep Learning. Aging Dis. 2021 Aug 1;12(5):1252–1262. doi: 10.14336/AD.2020.1202. PMID: 34341706; PMCID: PMC8279523.

[9] Dec E, Clement J, Cheng K, Church GM, Fossel MB, Rehkopf DH, Rosero-Bixby L, Kobor MS, Lin DT, Lu AT, Fei Z, Guo W, Chew YC, Yang X, Putra SED, Reiner AP, Correa A, Vilalta A, Pirazzini C, Passarino G, Monti D, Arosio B, Garagnani P, Franceschi C, Horvath S. Centenarian clocks: epigenetic clocks for validating claims of exceptional longevity. Geroscience. 2023 Jun;45(3):1817–1835. doi: 10.1007/s11357-023-00731-7. Epub 2023 Mar 25. PMID: 36964402; PMCID: PMC10400760.

[10] Snir S, Farrell C, Pellegrini M. Human epigenetic ageing is logarithmic with time across the entire lifespan. Epigenetics. 2019 Sep;14(9):912–926. doi: 10.1080/15592294.2019.1623634. Epub 2019 Jun 6. PMID: 31138013; PMCID: PMC6691990.

[11] Mboning L, Rubbi L, Thompson M, Bouchard LS, Pellegrini M. BayesAge: A maximum likelihood algorithm to predict epigenetic age. Front Bioinform. 2024 Apr 4;4:1329144. doi: 10.3389/fbinf.2024.1329144. PMID: 38638123; PMCID: PMC11024280.

[12] Mboning L, Costa EK, Chen J, Bouchard LS, Pellegrini M. BayesAge 2.0: A Maximum Likelihood Algorithm to Predict Transcriptomic Age. bioRxiv. 2024.09.16.613354. doi: 10.1101/2024.09.16.613354

[13] Morris TJ, Beck S. Analysis pipelines and packages for Infinium HumanMethylation450 BeadChip (450k) data. Methods. 2015 Jan 15;72:3–8. doi: 10.1016/j.ymeth.2014.08.011. Epub 2014 Sep 16. PMID: 25233806; PMCID: PMC4304832.

[14] Varshavsky M, Harari G, Glaser B, Dor Y, Shemer R, Kaplan T. Accurate age prediction from blood using a small set of DNA methylation sites and a cohort-based machine learning algorithm. Cell Rep Methods. 2023 Sep 25;3(9):100567. doi: 10.1016/j.crmeth.2023.100567. Epub 2023 Aug 28. PMID: 37751697; PMCID: PMC10545910.

[15] Hannum G, et al. Genome-wide methylation profiles reveal quantitative views of human aging rates. Mol Cell. 2013 Jan 24;49(2):359–367. doi: 10.1016/j.molcel.2012.10.016. Epub 2012 Nov 21. PMID: 23177740; PMCID: PMC3780611.

[16] van Dijk, S., Peters, T., Buckley, M. et al. DNA methylation in blood from neonatal screening cards and the association with BMI and insulin sensitivity in early childhood. Int J Obes 42, 28–35 (2018). 10.1038/ijo.2017.228

[17] Hannon E, et al. Characterizing genetic and environmental influences on variable DNA methylation using monozygotic and dizygotic twins. PLoS Genet. 2018 Aug 9;14(8):e1007544. doi: 10.1371/journal.pgen.1007544. PMID: 30091980; PMCID: PMC6084815.

[18] Hannon E, et al. DNA methylation meta-analysis reveals cellular alterations in psychosis and markers of treatment-resistant schizophrenia. Elife. 2021 Feb 26;10:e58430. doi: 10.7554/eLife.58430. PMID: 33646943; PMCID: PMC8009672.

[19] Kandaswamy, R., Hannon, E., et al. DNA methylation signatures of adolescent victimization: analysis of a longitudinal monozygotic twin sample. Epigenetics, 16(11), 1169–1186 (2020). 10.1080/15592294.2020.1853317

[20] Kho M, et al. Epigenetic loci for blood pressure are associated with hypertensive target organ damage in older African Americans from the genetic epidemiology network of Arteriopathy (GENOA) study. BMC Med Genomics. 2020 Sep 11;13(1):131. doi: 10.1186/s12920-020-00791-0. PMID: 32917208; PMCID: PMC7488710.

[21] Simo-Riudalbas L, et al. Genome-Wide DNA Methylation Analysis Identifies Novel Hypomethylated Non-Pericentromeric Genes with Potential Clinical Implications in ICF Syndrome. PLoS One. 2015 Jul 10;10(7):e0132517. doi: 10.1371/journal.pone.0132517. PMID: 26161907; PMCID: PMC4498748.

[22] Alisch RS, Barwick BG, et al. Age-associated DNA methylation in pediatric populations. Genome Res. 2012 Apr;22(4):623–32. doi: 10.1101/gr.125187.111. Epub 2012 Feb 1. PMID: 22300631; PMCID: PMC3317145.

[23] Horvath S, et al. Aging effects on DNA methylation modules in human brain and blood tissue. Genome Biol. 2012 Oct 3;13(10):R97. doi: 10.1186/gb-2012-13-10-r97. PMID: 23034122; PMCID: PMC4053733.

[24] Liu Y, et al. Epigenome-wide association data implicate DNA methylation as an intermediary of genetic risk in rheumatoid arthritis. Nat Biotechnol. 2013 Feb;31(2):142–7. doi: 10.1038/nbt.2487. Epub 2013 Jan 20. PMID: 23334450; PMCID: PMC3598632.

[25] Lehne B, et al. A coherent approach for analysis of the Illumina HumanMethylation450 BeadChip improves data quality and performance in epigenome-wide association studies. Genome Biol. 2015 Feb 15;16(1):37. doi: 10.1186/s13059-015-0600-x. Erratum in: Genome Biol. 2016 Apr 21;17:73. doi: 10.1186/s13059-016-0934-z. PMID: 25853392; PMCID: PMC4365767.

[26] Walker RF, et al. Epigenetic age analysis of children who seem to evade aging. Aging (Albany NY). 2015 May;7(5):334–9. doi: 10.18632/aging.100744. PMID: 25991677; PMCID: PMC4468314.

[27] Kananen L, et al. Aging-associated DNA methylation changes in middle-aged individuals: the Young Finns study. BMC Genomics. 2016 Feb 9;17:103. doi: 10.1186/s12864-016-2421-z. PMID: 26861258; PMCID: PMC4746895.

[28] Riccardi VM. Neurofibromatosis: The Importance of Localized or Otherwise Atypical Forms. Arch Dermatol. 1987;123(7):882–883. doi:10.1001/archderm.1987.01660310050011

[29] Horvath, S., Gurven, M., Levine, M.E. et al. An epigenetic clock analysis of race/ethnicity, sex, and coronary heart disease. Genome Biol 17, 171 (2016). 10.1186/s13059-016-1030-0

[30] Zannas AS, et al. Epigenetic upregulation of FKBP5 by aging and stress contributes to NF-κB-driven inflammation and cardiovascular risk. Proc Natl Acad Sci U S A. 2019 Jun 4;116(23):11370–11379. doi: 10.1073/pnas.1816847116. Epub 2019 May 21. PMID: 31113877; PMCID: PMC6561294.

[31] Voisin S, Almén MS, et al. Many obesity-associated SNPs strongly associate with DNA methylation changes at proximal promoters and enhancers. Genome Med. 2015 Oct 8;7:103. doi: 10.1186/s13073-015-0225-4. PMID: 26449484; PMCID: PMC4599317.

[32] Hannon, E., Dempster, E., Viana, J. et al. An integrated genetic-epigenetic analysis of schizophrenia: evidence for co-localization of genetic associations and differential DNA methylation. Genome Biol 17, 176 (2016). 10.1186/s13059-016-1041-x

[33] Ventham NT, et al. Integrative epigenome-wide analysis demonstrates that DNA methylation may mediate genetic risk in inflammatory bowel disease. Nat Commun. 2016 Nov 25;7:13507. doi: 10.1038/ncomms13507. PMID: 27886173; PMCID: PMC5133631.

[34] Heiss, J. A., Brennan, K. J., Baccarelli, A. A., Téllez-Rojo, M. M., Estrada-Gutiérrez, G., Wright, R. O., and Just, A. C. (2019). Battle of epigenetic proportions: comparing Illumina’s EPIC methylation microarrays and TruSeq targeted bisulfite sequencing. Epigenetics, 15(1–2), 174–182. 10.1080/15592294.2019.1656159

[35] Sugden K, et al. Patterns of Reliability: Assessing the Reproducibility and Integrity of DNA Methylation Measurement. Patterns (N Y). 2020 May 8;1(2):100014. doi: 10.1016/j.patter.2020.100014. Epub 2020 Apr 23. PMID: 32885222; PMCID: PMC7467214.

[36] Pauli Virtanen, et al. (2020) SciPy 1.0: Fundamental Algorithms for Scientific Computing in Python. Nature Methods, 17(3), 261–272.

[37] Seabold, Skipper, and Josef Perktold. “statsmodels: Econometric and statistical modeling with python.” Proceedings of the 9th Python in Science Conference. 2010.

[38] Oblak L, van der Zaag J, Higgins-Chen AT, Levine ME, Boks MP. A systematic review of biological, social and environmental factors associated with epigenetic clock acceleration. Ageing Res Rev. 2021 Aug;69:101348. doi: 10.1016/j.arr.2021.101348. Epub 2021 Apr 28. PMID: 33930583.

[39] Hui Zou, Trevor Hastie, Regularization and Variable Selection Via the Elastic Net, Journal of the Royal Statistical Society Series B: Statistical Methodology, Volume 67, Issue 2, April 2005, Pages 301–320, 10.1111/j.1467-9868.2005.00503.x

