## Supplementary material for "MicroBayesAge: A Maximum Likelihood Approach to Predict Epigenetic Age Using Microarray Data": MicroBayesAge Supplemental

NICOLE NOLAN

October 2024

(a) All male patients

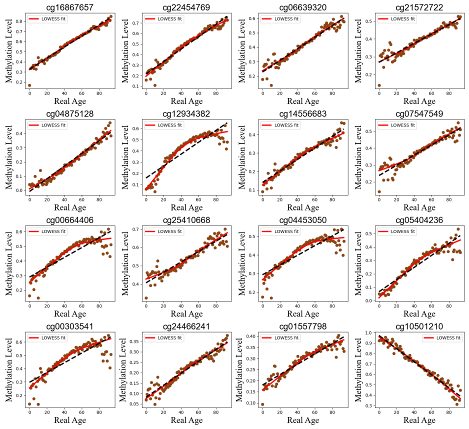


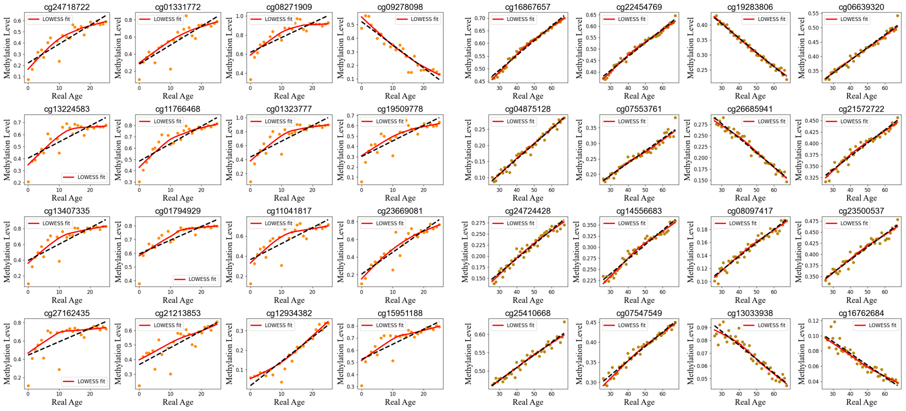


(b) Young male patients (c) Old male patients

Figure 1: Comparison of LOWESS regression fits with tau of 0.7 of the relationship between methylation and age for the top 16 most correlated CpG sites for all male patients, young male patients only, and old male patients only.

(a) All female patients

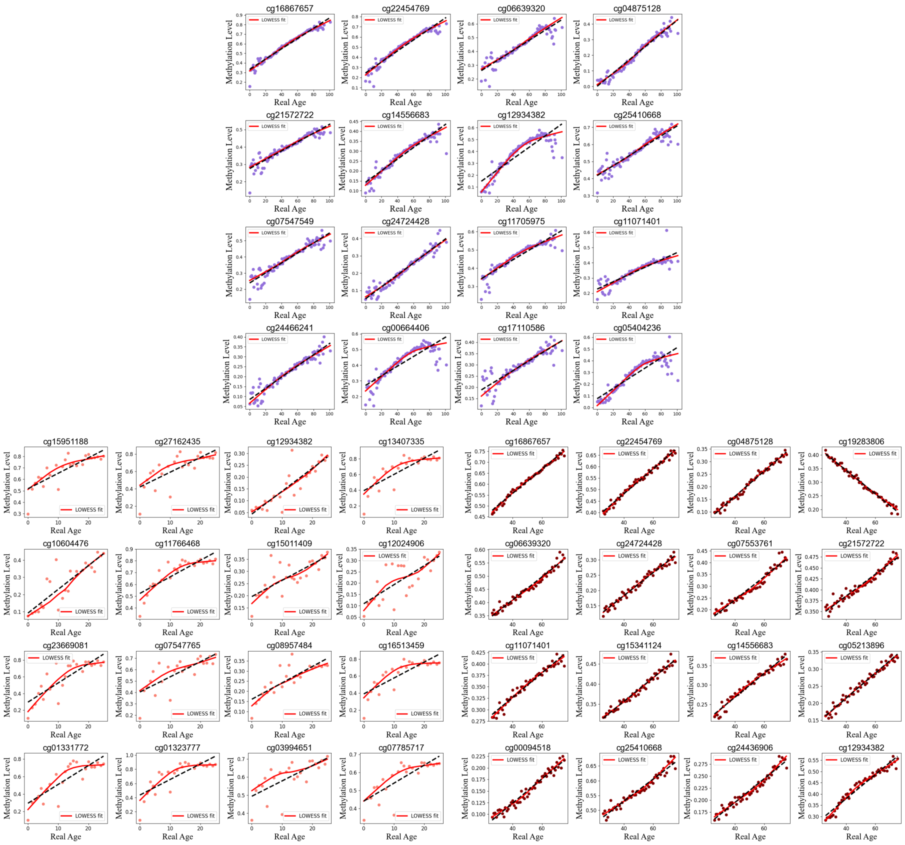


(b) Young female patients

(c) Old female patients

Figure 2: Comparison of LOWESS regression fits with tau of 0.7 of the relationship between methylation and age for the top 16 most correlated CpG sites for all female patients, young female patients only, and old female patients only.

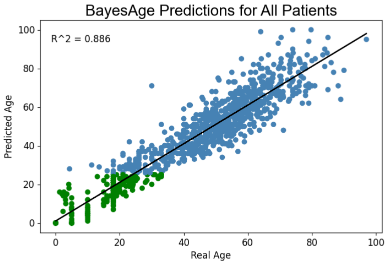


(a) First Stage Predictions

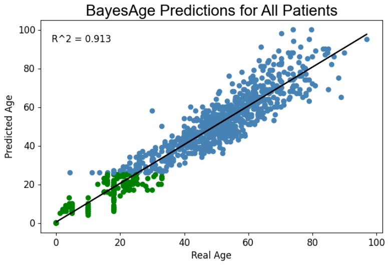


(b) Second Stage Predictions

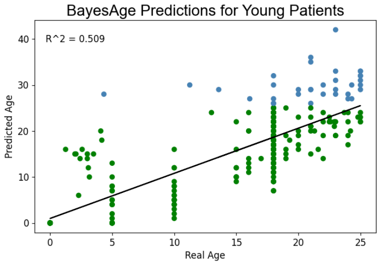


(c) First Stage Predictions

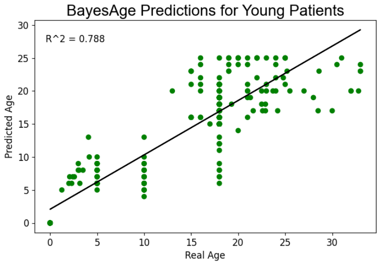


(d) Second Stage Predictions

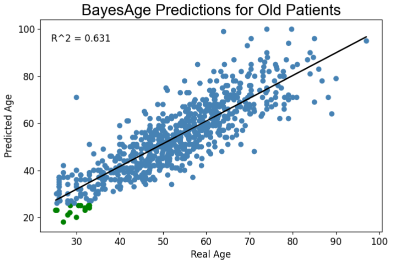


(e) First Stage Predictions

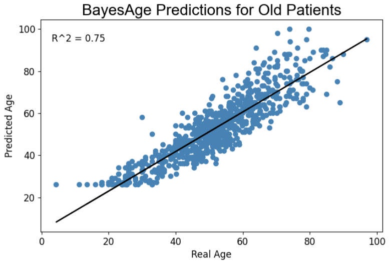


(f) Second Stage Predictions

Figure 3: Comparison of first and second stage age predictions plotted against real age for each age group of patients. Patients with predicted ages older than 25 are shown in blue while patients with predicted ages of 25 or younger are shown in green.

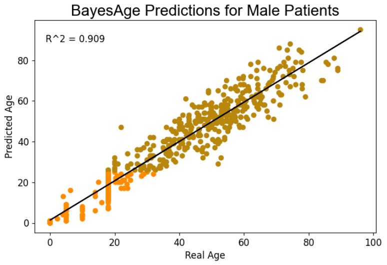


(a) First Stage Predictions

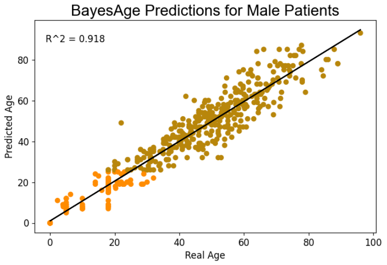


(b) Second Stage Predictions

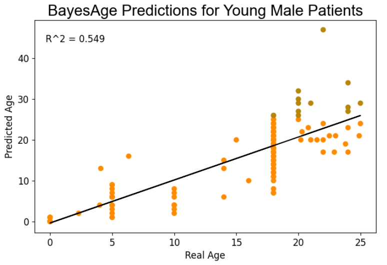


(c) First Stage Predictions

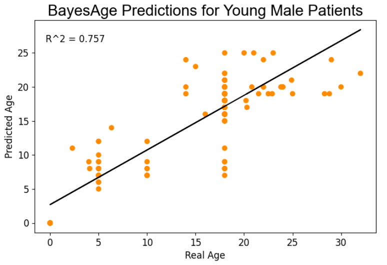


(d) Second Stage Predictions

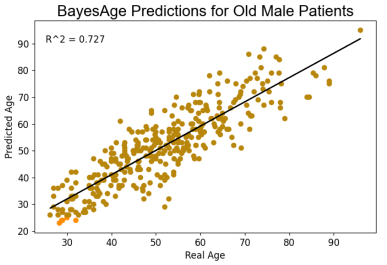


(e) First Stage Predictions

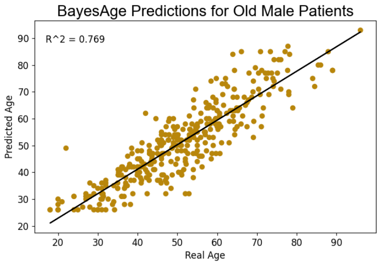


(f) Second Stage Predictions

Figure 4: Comparison of first and second stage age predictions plotted against real age for each age group of male patients. Male patients with predicted ages older than 25 are shown in yellow while patients with predicted ages of 25 or younger are shown in orange.

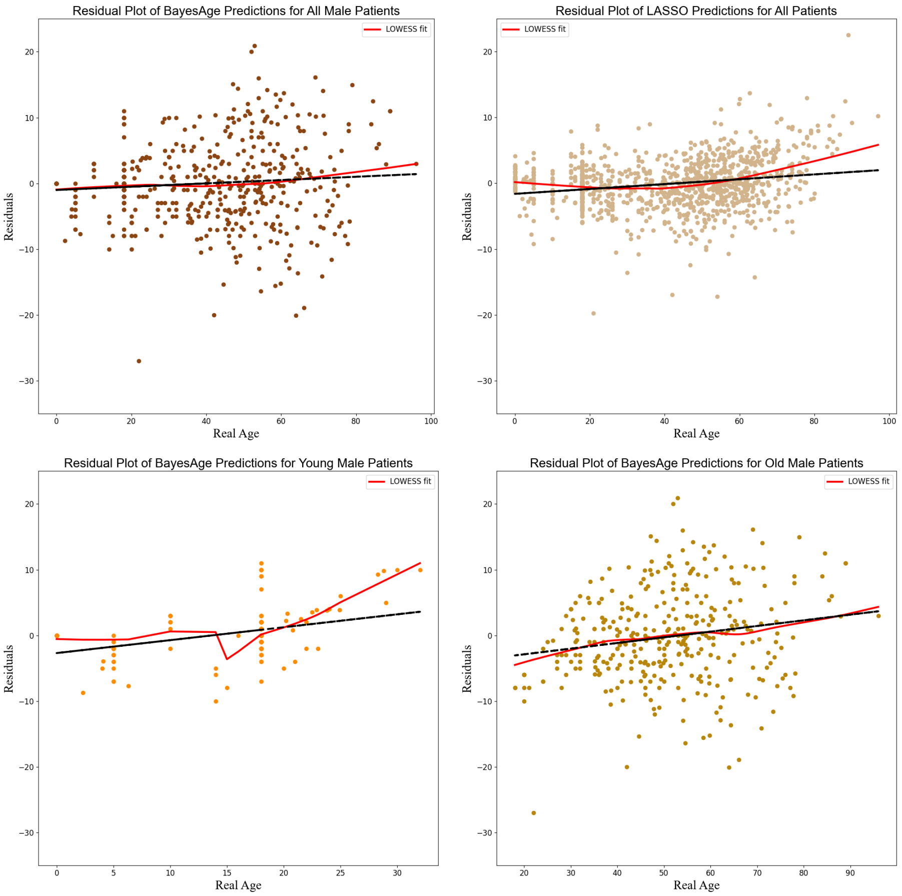


Figure 5: Residual plot of age predictions for male patients. First stage age predictions for male patients are shown in brown. Second stage age predictions are shown in yellow for older male patients and in orange for younger male patients. LASSO age predictions are shown in tan.

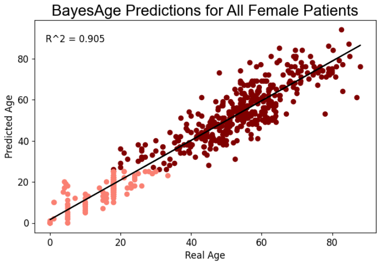


(a) First Stage Predictions

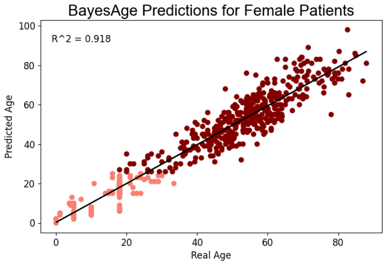


(b) Second Stage Predictions

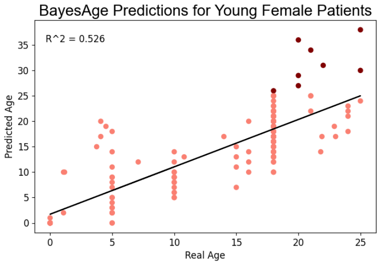


(c) First Stage Predictions

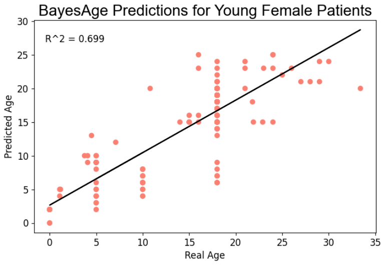


(d) Second Stage Predictions

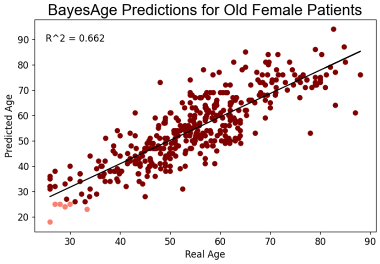


(e) First Stage Predictions

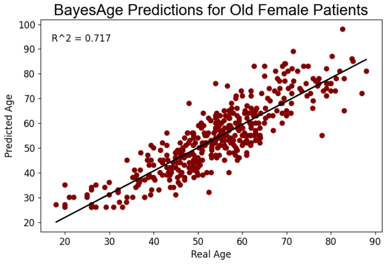


(f) Second Stage Predictions

Figure 6: Comparison of first and second stage age predictions plotted against real age for each age group of female patients. Female patients with predicted ages older than 25 are shown in red while patients with predicted ages of 25 or younger are shown in pink.

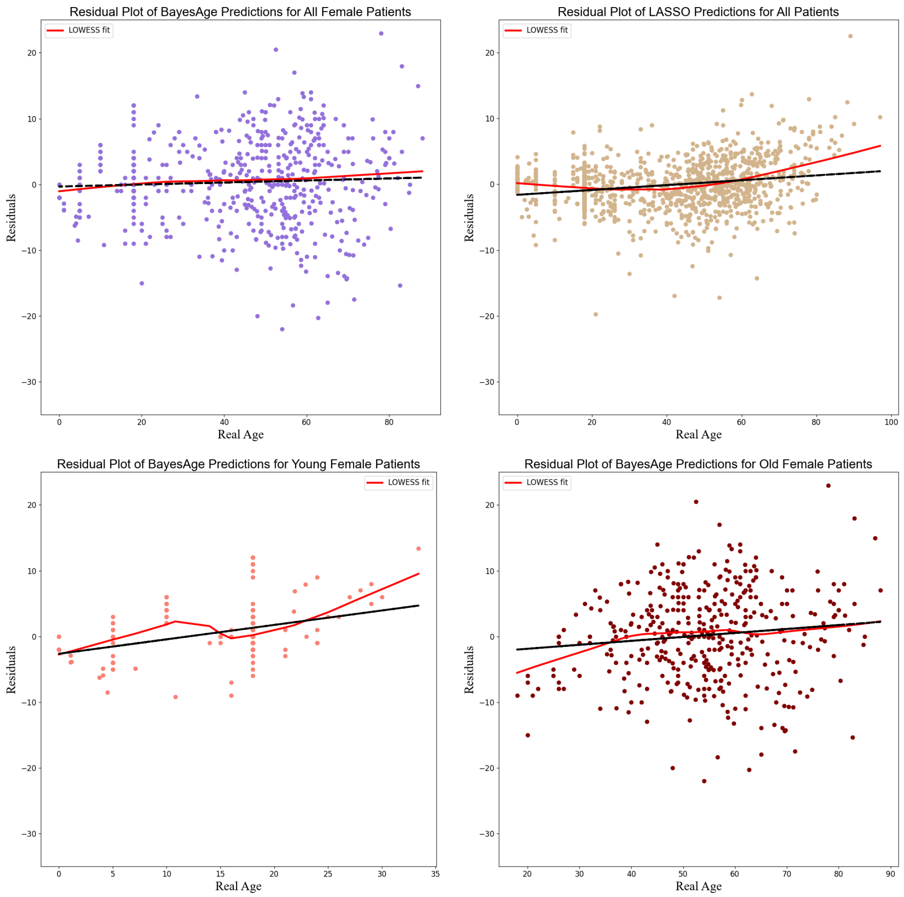


Figure 7: Residual plot of age predictions for female patients. First stage age predictions for female patients are shown in purple. Second stage age predictions are shown in red for older female patients and in pink for younger female patients. LASSO age predictions are shown in tan.
